# Plant-adapted OTO staining improves ultrastructural imaging by transmission electron microscopy

**DOI:** 10.64898/2026.09.08.750243

**Authors:** Huiping Yao, Xianhao Jin, Mengsi Sun, Haijing Cheng, Yuke Liao, Yantong Jiang, Huijuan Zhao, Hua Zhou

## Abstract

Transmission electron microscopy (TEM) is widely used to examine plant cellular ultrastructure, but sample preparation remains challenging because polysaccharide-rich cell walls, large vacuoles and complex membrane systems compromise staining and structural preservation. Here, we developed a plant-adapted TEM preparation method based on a modified osmium-thiocarbohydrazide-osmium (OTO) staining workflow combined with optimized washing, dehydration and resin infiltration. Using tomato roots and leaves, we show that the optimized method produces cleaner backgrounds, better-defined cellular boundaries and improved preservation of cellular morphology compared with conventional preparation. Fine membrane-associated structures, including mitochondrial cristae, chloroplast grana and stroma lamellae, nuclear membranes and other endomembrane structures, were more clearly resolved, while staining-related precipitates and diffuse background artifacts were reduced. This method provides a practical approach for high-contrast TEM imaging of plant tissues and ultrastructural analysis of plant cells and organelles.

---

Dear Editor,

Ultrastructural imaging provides essential information on the organization of plant cells and organelles that cannot be resolved by conventional light microscopy (Wang et al. 2019). Electron microscopy (EM), particularly TEM, has therefore played a central role in revealing the architecture of chloroplasts, mitochondria, endomembrane systems and other subcellular structures in plants (Otegui and Pennington 2019; Weiner et al. 2022). The quality and interpretability of electron micrographs, however, depend strongly on specimen preparation, including fixation, heavy-metal staining, dehydration, resin infiltration and sectioning (Wu et al. 2012; Wilson and Bacic 2012; Peddie et al. 2022).

Plant tissues present particular challenges for EM preparation. Rigid cell walls, large central vacuoles, intercellular air spaces and, in some tissues, waxy surface layers can impede reagent penetration and complicate uniform fixation, staining and resin infiltration(Wickramanayake and Czymmek 2023). In addition, membrane-rich organelles frequently occur adjacent to highly vacuolated or polysaccharide-rich cellular regions, creating a substantial dynamic range of electron density within the same section. Obtaining sufficient membrane contrast without generating excessive background or obscuring fine structural details is therefore a recurring challenge in plant TEM. The osmium–thiocarbohydrazide–osmium (OTO) approach was originally introduced to enhance the contrast of osmium-bound, lipid-rich cellular structures through thiocarbohydrazide (TCH)-mediated amplification of osmium deposition (Seligman et al. 1966; Willingham and Rutherford 1984; Hua et al. 2015). OTO-derived en bloc staining schemes were subsequently developed to generate strong and relatively homogeneous heavy-metal contrast for volume electron microscopy, particularly in neuronal tissues (Tapia et al. 2012; Mikula and Denk 2015; Hua et al. 2015; Song et al. 2023). More recently, modified OTO protocols have been applied to plant volume EM, where cell walls, vacuoles and tissue heterogeneity impose distinct preparation constraints (Kit^TEL^mann et al. 2016; Czymmek et al. 2020; Wickramanayake and Czymmek 2023). Lead aspartate staining provides an additional means of enhancing en bloc electron contrast and can be performed at elevated temperature to improve staining efficiency (Walton 1979; Tapia et al. 2012). Together, these developments demonstrate the power of sequential heavy-metal staining, but they also highlight an important challenge: maximizing heavy-metal deposition does not necessarily maximize ultrastructural information.

Many established OTO-based en bloc staining strategies were developed and optimized for animal tissues and rely on sequential or prolonged heavy-metal treatments to achieve high electron density. When transferred directly to plant material, however, increasing staining intensity alone did not consistently improve ultrastructural visualization. Excessive heavy-metal deposition has been reported to increase nonspecific electron-dense background and obscure fine membrane-associated structures, thereby compromising ultrastructural interpretability (Hart et al. 2024). We therefore optimize the balance between contrast generation and background removal rather than simply increasing stain concentration or incubation time. To achieve this, we developed a plant-adapted OTO workflow that integrates controlled sequential heavy-metal staining with stringent reagent preparation, segmented washing and staged resin infiltratio (Figure 1a). The protocol retains the overall framework of conventional resin-embedded TEM preparation while introducing coordinated modifications throughout the staining and sample-processing workflow (Table 1). Tomato (*Solanum lycopersicum*) roots and leaves were used to evaluate its performance across tissues with contrasting cellular architectures. Fresh tissues were first cut into small blocks (≤1 mm^3^) to facilitate rapid fixation and reagent penetration and were fixed with glutaraldehyde. Samples were then subjected to low-temperature ferrocyanide-reduced osmium staining, followed by TCH treatment and a second osmium staining step. Uranyl acetate staining and temperature-controlled Walton’s lead aspartate staining were subsequently incorporated to enhance electron contrast. Rather than relying on extended exposure to individual stains, we placed particular emphasis on controlling the transition between staining steps. Rapid solution exchange was followed by prolonged agitation-based washes to remove unbound or residual reagents before subsequent heavy-metal treatment. TCH was freshly prepared, fully dissolved and filtered before use to minimize crystalline contamination, while Walton’s lead staining was performed under controlled temperature and pH conditions, followed by temperature-matched washing. Gradient dehydration, progressive resin infiltration and extended polymerization were further incorporated to improve solvent exchange, embedding uniformity and section integrity (Figure 1a; Table 1). This combination proved important because artifacts generated at one stage of preparation frequently propagated through subsequent steps. Residual TCH or heavy-metal reagents, for example, could become sources of granular or crystalline deposits after later staining, whereas incomplete dehydration or resin infiltration could compromise section integrity despite otherwise satisfactory fixation and staining. The optimized protocol therefore treats staining, washing, dehydration and embedding as an integrated workflow rather than as independent procedures.

**Table 1.**
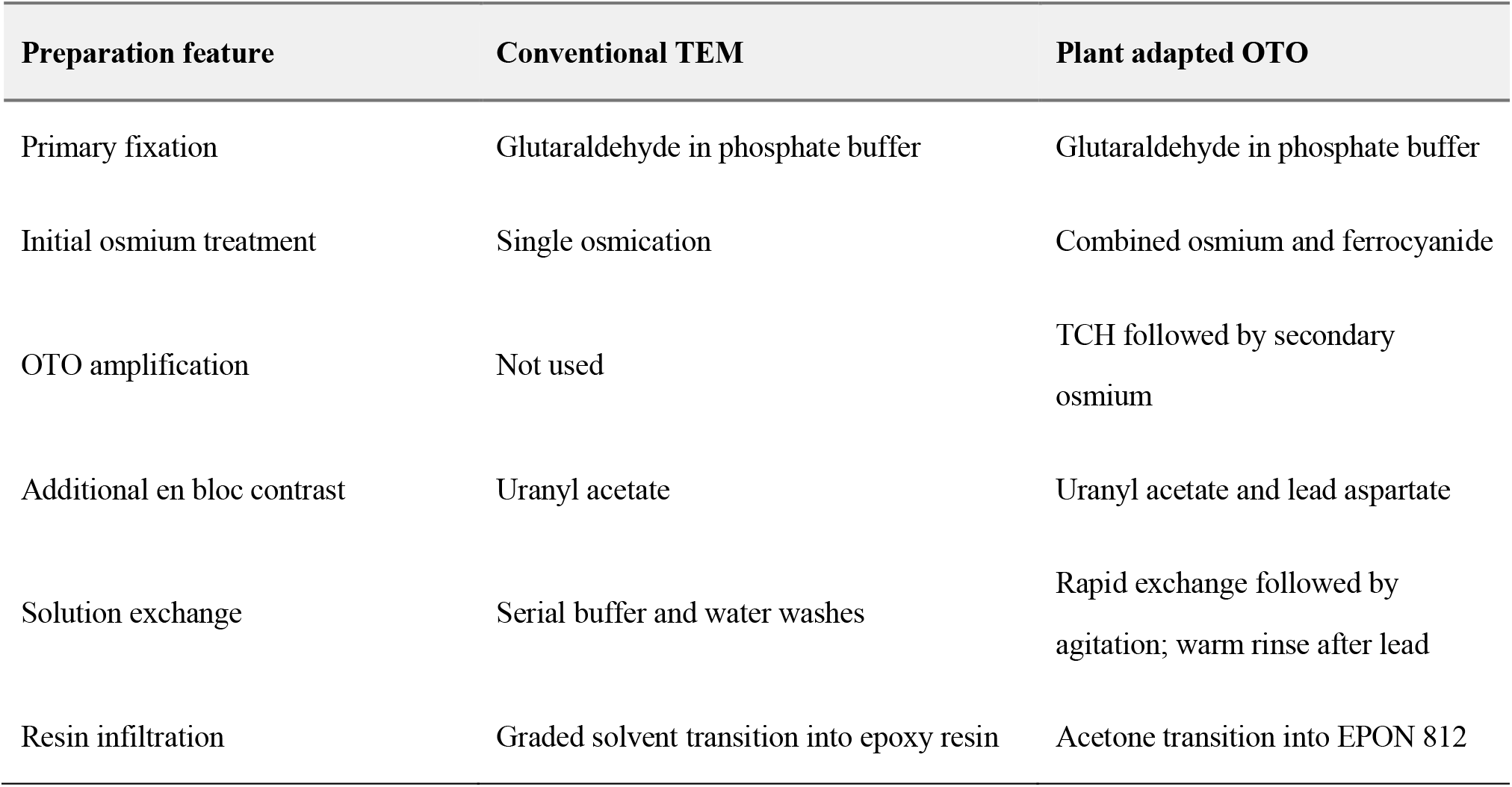
Comparison of conventional TEM preparation and the plant-adapted OTO workflow.

| Preparation feature | Conventional TEM | Plant adapted OTO |
| --- | --- | --- |
| Primary fixation | Glutaraldehyde in phosphate buffer | Glutaraldehyde in phosphate buffer |
| Initial osmium treatment | Single osmication | Combined osmium and ferrocyanide |
| OTO amplification | Not used | TCH followed by secondary osmium |
| Additional en bloc contrast | Uranyl acetate | Uranyl acetate and lead aspartate |
| Solution exchange | Serial buffer and water washes | Rapid exchange followed by agitation; warm rinse after lead |
| Resin infiltration | Graded solvent transition into epoxy resin | Acetone transition into EPON 812 |

**Figure 1.**
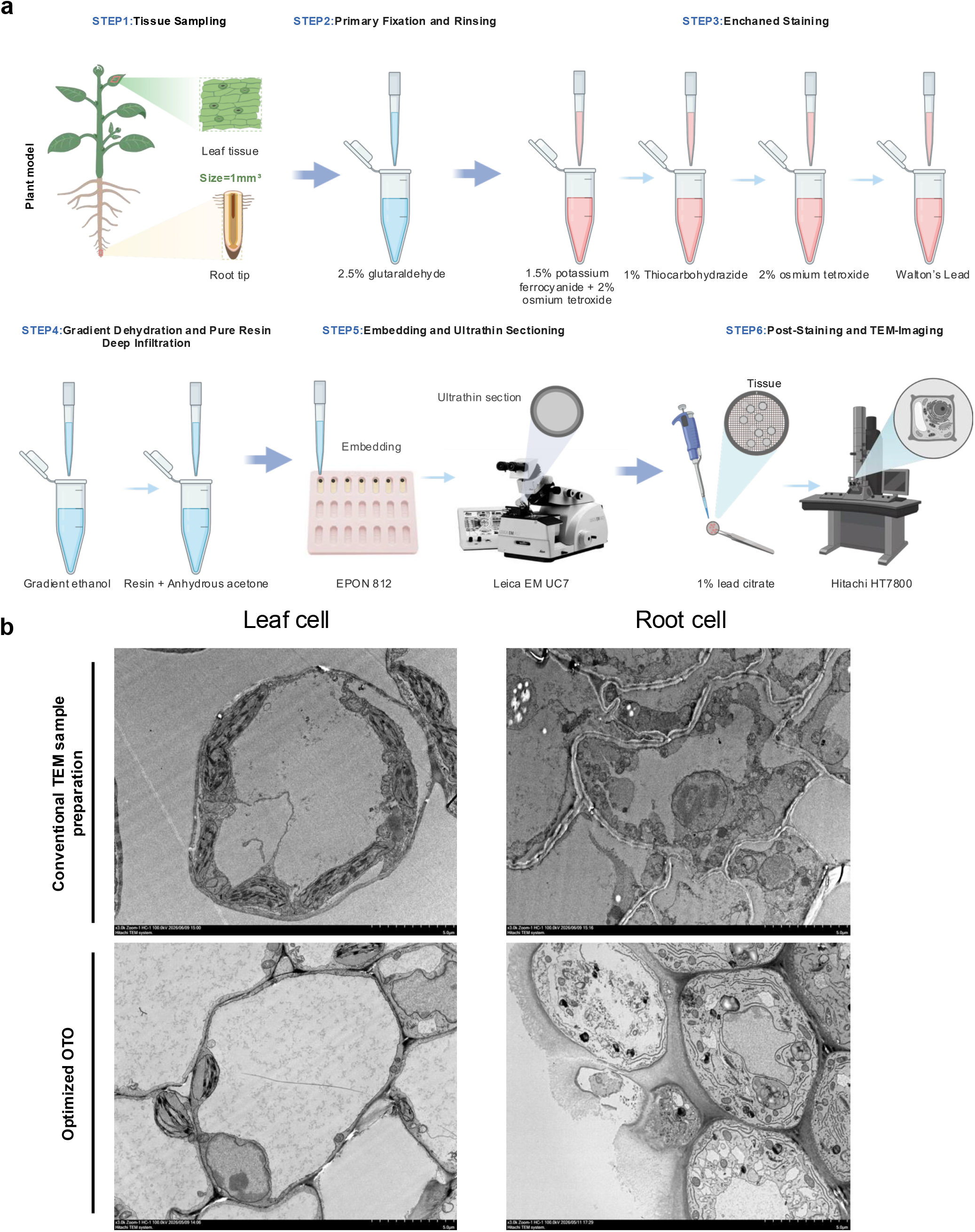
Overall performance of the optimized plant-adapted OTO workflow. **a**, Schematic overview of the optimized TEM preparation procedure, including fixation, ferrocyanide-reduced osmium staining, TCH-mediated enhancement, secondary osmium staining, uranyl acetate staining, Walton’s lead staining, segmented washing, gradient dehydration, resin infiltration, embedding and ultrathin sectioning. Key experimental conditions are indicated. **b**, Representative TEM images comparing conventionally prepared and optimized tomato tissues. The optimized workflow yields more uniform contrast, cleaner backgrounds and better-defined cellular boundaries. Scale bars are indicated in each panel.

Compared with conventional preparation, the optimized workflow produced more homogeneous cellular contrast and substantially cleaner backgrounds (Figure 1b). Cell contours remained well preserved, and membrane-associated boundaries were more readily distinguished from the surrounding cytoplasm. The improvement was particularly apparent in regions in which conventionally prepared samples displayed diffuse electron-dense background or spatially uneven staining. Optimized samples also contained fewer scattered electron-dense precipitates, reducing structures that could otherwise be mistaken for genuine cellular features. Common artifacts encountered further demonstrated the importance of reagent preparation and washing. Incompletely dissolved or unfiltered staining solutions generated granular or crystalline deposits, whereas insufficient washing between successive heavy-metal treatments resulted in diffuse background accumulation. Deviations from appropriate TCH preparation or from the temperature and pH conditions required for lead aspartate staining similarly compromised image clarity. In addition, incomplete dehydration and resin infiltration contributed to fractured sections and localized loss of structural integrity. Fresh reagent preparation, filtration, sequential solution exchange and controlled dehydration and infiltration substantially reduced these artifacts. Thus, successful implementation of OTO staining in plant tissues depends not only on the amount of heavy metal deposited, but also on controlling where staining reagents accumulate and how efficiently excess reagent is removed.

We next asked whether the improved overall image quality translated into better visualization of cellular and subcellular structures in leaf tissue. Leaf mesophyll cells provide a useful system for evaluating TEM preparation because they contain large vacuoles together with abundant chloroplasts, mitochondria and other membrane-rich organelles, allowing structural preservation and contrast to be assessed at multiple scales within the same tissue. At the cellular scale, conventionally prepared samples frequently showed a diffuse grey background and poorly resolved intracellular boundaries, which reduced the distinction between the cell wall, cytoplasm and vacuolar compartment (Figure 2a). In samples processed using the optimized workflow, the cytoplasmic background was more homogeneous, cellular boundaries were more clearly defined and intracellular structures could be distinguished more readily. The improved contrast therefore enhanced the overall interpretability of mesophyll cell organization without obscuring fine structural details. The advantages of the optimized preparation became more apparent at higher magnification. Chloroplasts are particularly demanding structures for evaluating TEM preparation because their closely apposed thylakoid membranes require strong membrane contrast while retaining sufficient separation to resolve individual lamellae. In conventionally prepared samples, diffuse staining frequently reduced the distinction between adjacent thylakoid membranes, making individual grana stacks and connecting stroma lamellae difficult to resolve. In optimized samples, the chloroplast envelope was well preserved, while grana stacks and stroma lamellae appeared more continuous and clearly separated from the surrounding stroma (Figure 2b). Importantly, this improved structural definition was achieved without excessive electron density that could obscure the internal organization of the chloroplast. Mitochondrial ultrastructure similarly benefited from the optimized workflow. Conventional preparations often showed limited differentiation between the mitochondrial matrix and internal membranes, and individual cristae were not consistently resolved. In contrast, mitochondria prepared using the optimized protocol exhibited well-defined outer boundaries and more readily identifiable cristae (Figure 2c). These observations indicate that the optimized staining conditions provide sufficient membrane-associated contrast while maintaining the spatial definition of closely packed internal membranes. Improved structural differentiation was also evident in nuclei. Compared with conventional preparation, optimized samples showed a more clearly defined nuclear envelope and improved distinction between regions of differing electron density within the nucleoplasm (Figure 2d). Nuclear-envelopeassociated structures were also more readily identifiable where appropriately oriented sections were obtained.

**Figure 2.**
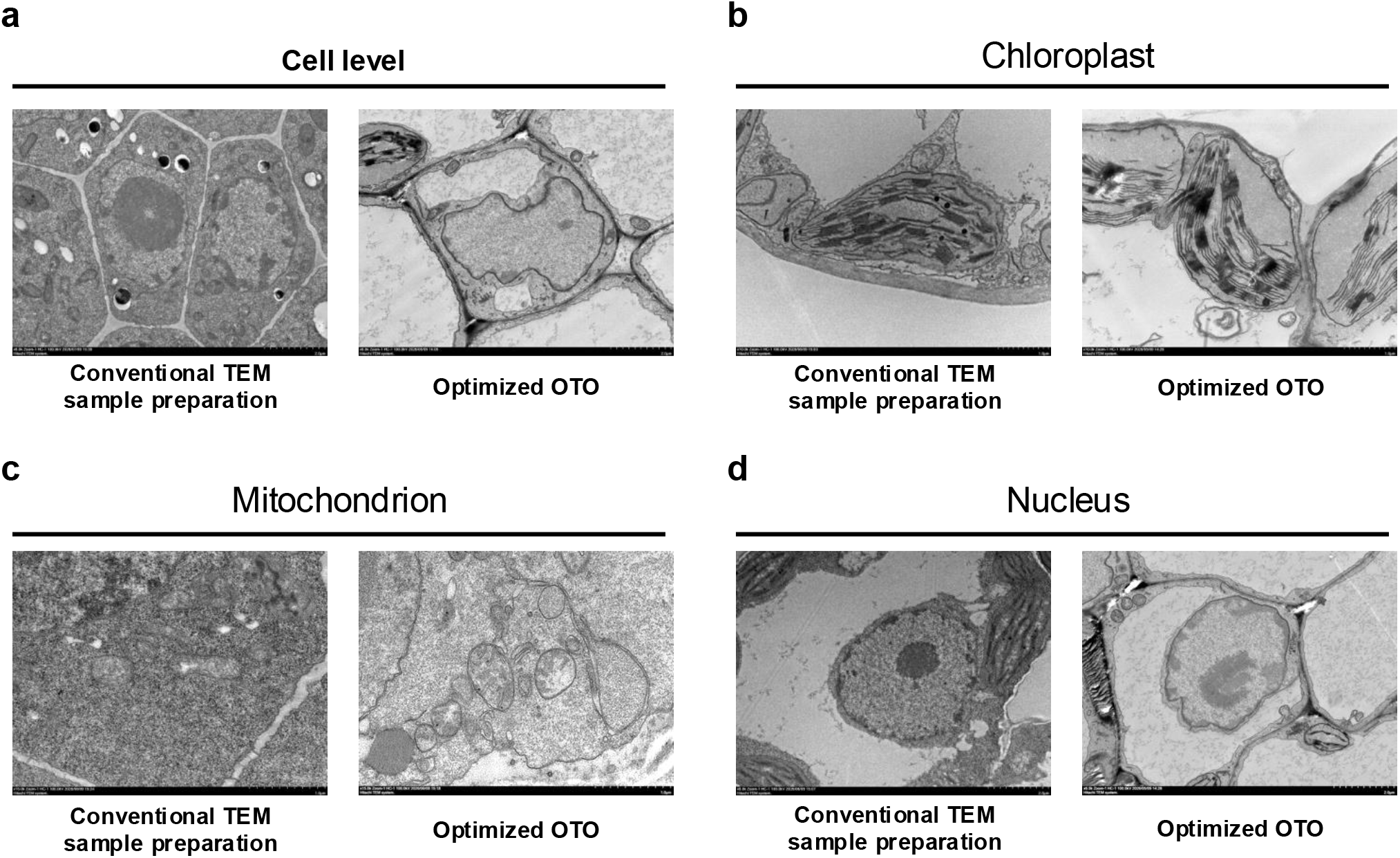
Enhanced visualization of cellular and organelle ultrastructure using the optimized plant-adapted OTO workflow. **a**, Representative leaf cells prepared using conventional and optimized methods, showing differences in background staining, cell-wall definition, membrane continuity and intracellular structural preservation. **b**, High-magnification comparison of chloroplast ultrastructure. The optimized preparation provides clearer visualization of the chloroplast envelope, grana stacks and stroma lamellae. **c**, Representative mitochondria showing improved definition of mitochondrial boundaries and cristae following optimized preparation. **d**, Nuclear ultrastructure, including the nuclear envelope, nuclear pores and chromatin organization, under conventional and optimized preparation conditions. Enlarged regions highlight representative structural differences. Scale bars are indicated in each panel.

A central feature of the workflow is the coordination of sequential staining, washing and embedding rather than reliance on any single staining step. Reduced osmium, TCH-mediated amplification, secondary osmium, uranyl acetate and heated lead aspartate together provide sufficient electron contrast, while extensive intermediate washing limits reagent carry-over and precipitation. Gradual dehydration and staged resin infiltration further preserve structural integrity. Thus, the improved image quality reflects a balance between contrast enhancement and avoidance of excessive or uneven heavy-metal deposition. Because the protocol uses conventional chemical fixation and resin embedding, it can be readily implemented in laboratories already equipped for routine TEM. Its consistent performance in tomato roots and leaves also suggests applicability across tissues with distinct cellular architectures, although further adjustment may be required for particularly dense, lignified or highly cuticularized material.

## Materials and Methods

### Plant materials

Tomato (*Solanum lycopersicum* L.) seedlings were grown on Murashige and Skoog (MS) medium for 15 days. Roots and leaves were collected from 15-day-old seedlings and processed immediately after harvest. Samples from three independently grown seedlings were used as biological replicates.

### Reagents and solution preparation

Phosphate buffer (PB; 0.1 M, pH 7.2) was prepared from disodium hydrogen phosphate and sodium dihydrogen phosphate. The primary fixative consisted of 2.5% (v/v) EM-grade glutaraldehyde prepared in 0.1 M PB. Reduced osmium solution was freshly prepared by combining 2% (w/v) osmium tetroxide (OsO4) and 1.5% (w/v) potassium ferrocyanide in 0.1 M PB. Thiocarbohydrazide (TCH) was prepared as a 1% (w/v) aqueous solution, dissolved at 60 °C with intermittent mixing for at least 30 min, cooled to room temperature and filtered through a 0.22-µm membrane before use. Uranyl acetate was prepared as a 2% (w/v) aqueous solution and filtered before staining. Walton’s lead aspartate solution was prepared by dissolving 0.04 g L-aspartic acid and 0.066 g lead nitrate in 10 ml distilled water at 60 °C, followed by adjustment to pH 5.4 with sodium hydroxide. EPON 812 embedding resin was prepared by mixing 36.08 g EPON 812, 10.85 g dodecenyl succinic anhydride (DDSA), 22.39 g nadic methyl anhydride (NMA) and 0.88 g 2,4,6-tris(dimethylaminomethyl)phenol (DMP-30).

### Optimized OTO-based preparation

Fresh tissues were cut into blocks ≤1 mm^3^ and immediately fixed in 2.5% glutaraldehyde in 0.1 M PB (pH 7.2) for 60–90 min at room temperature, followed by overnight fixation at 4 °C. Samples were washed thoroughly with 0.1 M PB. For the first heavy-metal staining step, samples were incubated for 1 h on ice in the dark in freshly prepared solution containing 1.5% potassium ferrocyanide and 2% OsO4 in 0.1 M PB. After extensive washing with water, samples were treated with freshly prepared 1% TCH for 40 min at room temperature and subsequently washed again. Samples were then incubated in 2% aqueous OsO4 for 60 min at room temperature in the dark, washed with water and stained with 2% aqueous uranyl acetate for 30 min at room temperature. After washing, samples were incubated in Walton’s lead aspartate solution at 60 °C for 30 min. Residual lead solution was removed by washing once with preheated water at 60 °C, followed by three washes with water at room temperature.

### Dehydration, resin infiltration and embedding

Samples were dehydrated at room temperature through 30%, 50%, 70%, 80% and 90% ethanol, followed by 90% ethanol:90% acetone (1:1, v/v) and 90% acetone, with 10 min at each step. Samples were then washed three times in 100% anhydrous acetone. Resin infiltration was performed sequentially using EPON 812:acetone mixtures of 1:3 for 2 h, 1:1 for 3 h and 3:1 for 4 h, followed by infiltration in 100% EPON 812 for 6–12 h. Pure resin was replaced three times before embedding. Samples were oriented in embedding moulds and polymerized at 60 °C for 48 h.

### Ultrathin sectioning and TEM imaging

Polymerized resin blocks were trimmed, and ultrathin sections of approximately 60 nm were cut using an ultramicrotome equipped with a diamond knife. Sections were collected on copper grids and examined using a Hitachi transmission electron microscope operated at an accelerating voltage of 100 kV. Digital images were acquired using the microscope imaging system. Corresponding tissues and subcellular structures from samples prepared using the conventional and optimized protocols were imaged at comparable magnifications and acquisition settings to allow direct comparison of ultrastructural preservation and contrast.

### Image assessment

TEM images were processed using Fiji. Directly compared images were adjusted using equivalent brightness and contrast settings. Method performance was assessed based on background uniformity, cellular and membrane boundary definition, structural preservation, staining-associated precipitates and visibility of organelle ultrastructure.

## Acknowledgments

This work was supported by Hangzhou Tsientang Education Foundation.

## Author Contributions

H.Y., X.J., M.S. and H.Z. conceived and designed the study. H.Y., X.J., M.S. and H.C., performed the experiments. H.Y., X.J., M.S. and H.C. analyzed the data. Y.L., Y.J. and H.Z. provided materials, technical support and conceptual insights. H.Z. supervised and oversaw the study. H.Y., X.J., M.S. and H.Z. wrote the manuscript, and all authors reviewed and revised the manuscript.

